# Growth hormone receptor (GHR) in AgRP neurons regulates thermogenesis in aged mice in a sex-specific manner

**DOI:** 10.1101/2022.08.31.506031

**Authors:** Lukas Stilgenbauer, Juliana Bezerra Medeiros de Lima, Lucas Kniess Debarba, Manal Khan, Lisa Koshko, John J. Kopchick, Andrzej Bartke, Augusto Schneider, Marianna Sadagurski

## Abstract

Evidence for hypothalamic regulation of energy homeostasis and thermoregulation in brown adipose tissue (BAT) during aging has been well recognized, yet the central molecular mediators involved in this process are poorly understood. The arcuate hypothalamus (ARC), orexigenic agouti-related peptide (AgRP) neurons control nutrient intake, energy homeostasis, and BAT thermogenesis. To determine the roles of growth hormone receptor (GHR) signaling in the AgRP neurons we used mice with the AgRP-specific GHR deletion (AgRP^ΔGHR^). We found that female AgRP^ΔGHR^ mice were resistant to temperature adaptation, and their body core temperature remained significantly lower when held at 10°C, 22°C, or 30°C, compared to control mice. Low body core temperature in female AgRP^ΔGHR^ mice has been associated with significant reductions in *Ucp1* and *Pgc1α* expression in the BAT. Further, neuronal activity in AgRP in response to cold exposure was blunted in AgRP^ΔGHR^ females, while the number of Fos^+^ AgRP neurons was increased in control females exposed to cold. Global transcriptome from BAT identified increased expression of genes related to immune responses and chemokine activity and decreased expression of genes involved in triglycerides synthesis and metabolic pathways in AgRP^ΔGHR^ females. Importantly, these were the same genes that are downregulated by thermoneutrality in control mice but not in the AgRP^ΔGHR^ animals. Collectively, these data demonstrate a novel circuit of thermal regulation between the hypothalamic AgRP-GHR and BAT and provide insight into the brain systems that are critical for the thermogenic vitality of the elderly.

## Introduction

During aging, BAT loses its thermogenic capacity and adaptation to cold temperatures thus reducing its ability to maintain normal energy homeostasis and body temperature late in life [1]. Energy homeostasis and thermoregulation are coordinated in the arcuate nucleus of the hypothalamus (ARC), which integrates neuronal and hormonal signals originating from peripheral tissues [2]. In the ARC, an antagonistic interaction between neurons expressing agouti-related peptide (AgRP) and neurons expressing proopiomelanocortin (POMC) constitutes the central metabolic controlling axis, and alterations in its activity impair energy homeostasis, thermoregulation, and peripheral glucose metabolism [3, 4]. Age-associated alterations in thermoregulation include low heat production, impaired thermal perception, and impaired autonomic and thermoregulatory responses [5]. Shivering is critical for heat production in a cold environment. Compared to young adults, shivering to increase heat production is impaired in elderly people and in laboratory animals, which reduces their tolerance for cold environments [6, 7].

In the hypothalamus, thermoregulatory circuits are modulated by leptin- and insulin-sensing neurons that respond to external nutrient and temperature cues [8]. Hypothalamic leptin and insulin signaling influence baselines of body core temperature in the fed state and oppose entry into a torpor-like state, without an effect on thermogenic response to cold [8]. In general, the preoptic area (POA) of the hypothalamus is responsible for managing body temperature, however, a subset of AgRP neurons was recently identified to mediate thermogenesis [9, 10]. In support, female mice lacking corticotropin-releasing factor receptor type 1 (CRFR1) selectively in AgRP neurons, exhibit a maladaptive thermogenic response to cold following impaired activation of the sympathetic nervous system (SNS) [11].

Growth hormone (GH) is a key mediator of growth and metabolism [12, 13]. GH deficient, long-lived Ames dwarf mice have highly active BAT compared to their littermates as indicated by depleted lipid stores, increased BAT tissue weight, increased expression of genes related to thermogenesis and lipid metabolism, and increased oxygen consumption and energy expenditure [14–16]. Housing Ames dwarf mice at thermoneutrality normalizes their lipid stores, gene expression, and oxygen consumption to those of control mice [14, 15], suggesting that BAT contributes to the extended longevity of GH-deficient mice.

GH activates AgRP neurons [17]. We and others have shown that young mice carrying AgRP-specific GHR ablation (AgRP^ΔGHR^ mice) have similar body weight, food intake, hormonal levels, and insulin sensitivity compared to control animals [17, 18]. However, during fasting AgRP neurons’ ability to save energy is impaired in AgRP GHR KO male mice, leading to increased fat loss, indicating GH as a starvation signal in AgRP neurons [17]. Given the role GH signaling plays in aging and its effect on BAT thermogenic capacity, in the current study we used AgRP^ΔGHR^ mice to explore the role of GHR signals in AgRP neurons in thermoregulation and thermoregulatory responses to temperature cues in aging animals.

## Materials and Methods

### Animals

Adult male and female AgRP^tm1(cre)^ (AgRP-Ires-cre, stock 012899) were purchased from The Jackson Laboratory, and GHR^L/L^ mice were described previously [19]. The characterization of the AgRP^ΔGHR^ mice was described previously [18]. We used wild-type littermate controls whenever possible and if not available, we used age-matched controls from the same breeding line. All mice were provided *ad libitum* access to a standard chow diet (Purina Lab Diet 5001) and housed in temperature-controlled rooms on a 12-hour/12-hour light-dark cycle. Procedures involved in this study were approved by the Wayne State University Institutional Animal Care and Use Committee (IACUC).

### Temperature Exposure and Core Body Temperature Monitoring

Mice were anesthetized using isoflurane. A mid-sagittal incision was made in the abdomen and a passive RFID chip was inserted into the abdominal fat pat. Mice were monitored three times daily with RFID readers for core body temperature. Mice were housed at 22°C for two days before the temperature challenge to allow them to equilibrate. After two days of equilibration, the mice were then monitored for two days at 22°C followed by 3 days at 10°C, two days at 22°C, and three days at 30°C. Metabolic measurements of energy homeostasis at 22°C were obtained using an indirect calorimetry system (PhenoMaster, TSE system, Bad Homburg, Germany). The mice were acclimatized to the cages for 2 days and monitored for 5 days while food and water were provided *ad libitum*.

### Perfusion and immunolabeling

Mice were anesthetized and perfused using phosphate buffer saline (PBS) (pH 7.5) followed by 4% paraformaldehyde (PFA). Brains were post-fixed, dehydrated, and sectioned coronally (30μm) using a sliding microtome, followed by immunofluorescent analysis as described [20]. For immunohistochemistry brain sections were washed with PBS six times, blocked with 0.3% Triton X-100 and 3% normal donkey serum in PBS for 2h; then the staining was carried out with the mouse anti-cFos (anti-rabbit, 1:500, cat. number sc-52, Santa Cruz). Immunostaining brain sections were pretreated with 0.5% NaOH and 0.5% H_2_O_2_ in PBS for 20 min. After the primary antibody, brain sections were incubated with AlexaFluor-conjugated secondary antibodies for 2h (Invitrogen). Microscopic images of the stained sections were obtained using an Olympus FluoView 500 and Laser Scanning Confocal Microscope Zeiss LSM 800.

### Quantification

For the quantification of immunoreactivity, images of matched brain areas were taken from at least 3 sections containing the hypothalamus for each brain between bregma −0.82 mm to −2.4 mm (according to the Franklin mouse brain atlas). Serial brain sections were made at 30 μm thickness, and every five sections were represented by one section with staining and cell counting. All sections were arranged from rostral to caudal to examine the distribution of labeled cells. cFos^+^ cells were counted using ImageJ with DAPI (nuclear staining). The average of the total number of cells/field was assessed by statistical analysis as detailed below.

### Quantitative Real-Time PCR

Total RNA was isolated from dissected BAT using Trizol reagent (Invitrogen, #15596026). The concentration of 1000 ng of RNA was used for cDNA synthesis using a High Capacity cDNA Reverse Transcription Kit (BioRad, #1708891). To detect the contaminated DNA we used the samples processed without the transcriptase reverse enzyme as negative controls. Quantitative real-time PCR was performed using the Applied Biosystems 7500 Real-Time PCR System. PGC1α, Forward: GCAACATGCTCAAGCCAAAC, Reverse: TGCAGTTCCAGAGAGTTCCA; Ucp1, Forward: GCTTTGCCTCACTCAGGATTGG, Reverse: CCAATGAACACTGCCACACCTC; FGF21, Forward: CCTCTAGGTTTCTTTGCCAACAG, Reverse: AAGCTGGCCTCAGGAT. Each PCR reaction was performed in duplicate. As negative controls, we used water instead of the cDNA, and ß-actin was measured in each cDNA sample as the housekeeping gene. The ΔΔCT method was used to determine the gene transcripts in each sample. For each sample, the threshold cycle (CT) was measured and normalized to the average of the housekeeping gene (ΔCT = CT gene of interest - CT housekeeping gene). The fold change of mRNA in the rest of the samples relative to the control male group was determined by 2-ΔΔCT. Data are shown as mRNA expression levels relative to the control males.

### Histology and Morphometric Analysis

Histological analysis was performed on BAT tissues isolated from the animals at room temperature or exposed to 10°C, or at 30°C as previously described [21]. Morphometric analysis of BAT was performed with NIH ImageJ software (http://rsb.info.nih.gov/ij/).

### RNA Extraction and mRNA sequencing

RNA-Seq was performed at the WSU Genome Sciences Core. All RNA-Seq data processing was performed as before [22]. Transcriptomic profile of individual BAT samples was performed using commercial RNA-sequencing kits (NEBNext mRNA Library Prep Master Mix and NEBNext Multiplex Oligos for Illumina, New England Biolabs, Ipswich, MA) and adapted according to previous descriptions [22]. All RNA-Seq data are available at the Sequence Read Archive (SRA) at NCBI under accession number PRJNA871915. The mapping of sequencing reads to the mouse transcriptome and mRNA abundance was performed as previously [22]. mRNAs were further processed for pathway analysis using the Generally Applicable Gene-set Enrichment (GAGE), for the enrichment of KEGG pathways and gene ontology (GO) terms (biological processes, molecular function, and cellular component).

### Statistical analysis

Statistical analyses for differentially expressed mRNAs were performed pairwise using EdgeR in the software R (3.2.2). Genes with a false discovery rate (FDR) < 0.05 and fold change (FC) > 2.0 were considered upregulated and with FDR < 0.05 and FC < 0.5 were considered downregulated. For all other experiments, the unpaired two-tailed Student’s *t*-test was used for comparisons between two groups. Statistical analyses were performed using the GraphPad Prism software. A *p*-value of less than 0.05 was considered statistically significant.

## Results

### Female AgRP^ΔGHR^ mice exhibit decreased body core temperature

AgRP^ΔGHR^ mice show no difference in body weight as compared to control littermates (Figure 1A). Although, by 14 months of age female AgRP^ΔGHR^ mice demonstrated a slight, but insignificant (p=0.056), gain in body weight (Figure 1A). By 12 weeks of age, female but not male AgRP^ΔGHR^ mice exhibited reduced body core temperature compared to controls (Figures 1B and C). The body core temperature was overall higher in female mice, consistent with previous reports [23]. Additionally, when placed in metabolic chambers, female AgRP^ΔGHR^ mice exhibited a reduction in heat production without differences in the respiratory exchange ratio (RER) or ambulatory activity levels (Figure 1D and Supplementary Figure 1). These data are in agreement with a previous report showing that young AgRP^ΔGHR^ male mice exhibited no differences in heat production, RER or activity levels (Figure 1E and Supplementary Figure 1) [17]. Gene expression analysis of BAT demonstrated reduced levels of *Ucp1* (mitochondrial uncoupling protein 1) and *Pgc1α* (Ppargc1α, peroxisome proliferator-activated receptor gamma, coactivator 1 alpha) in 3 months old female AgRP^ΔGHR^ mice compared to control littermates (Figure 1F). We did not detect differences in the expression levels of *Ucp1* and *Pgc1α* in male AgRP^ΔGHR^ mice compared to controls (Figure 1G).

**Figure 1.**
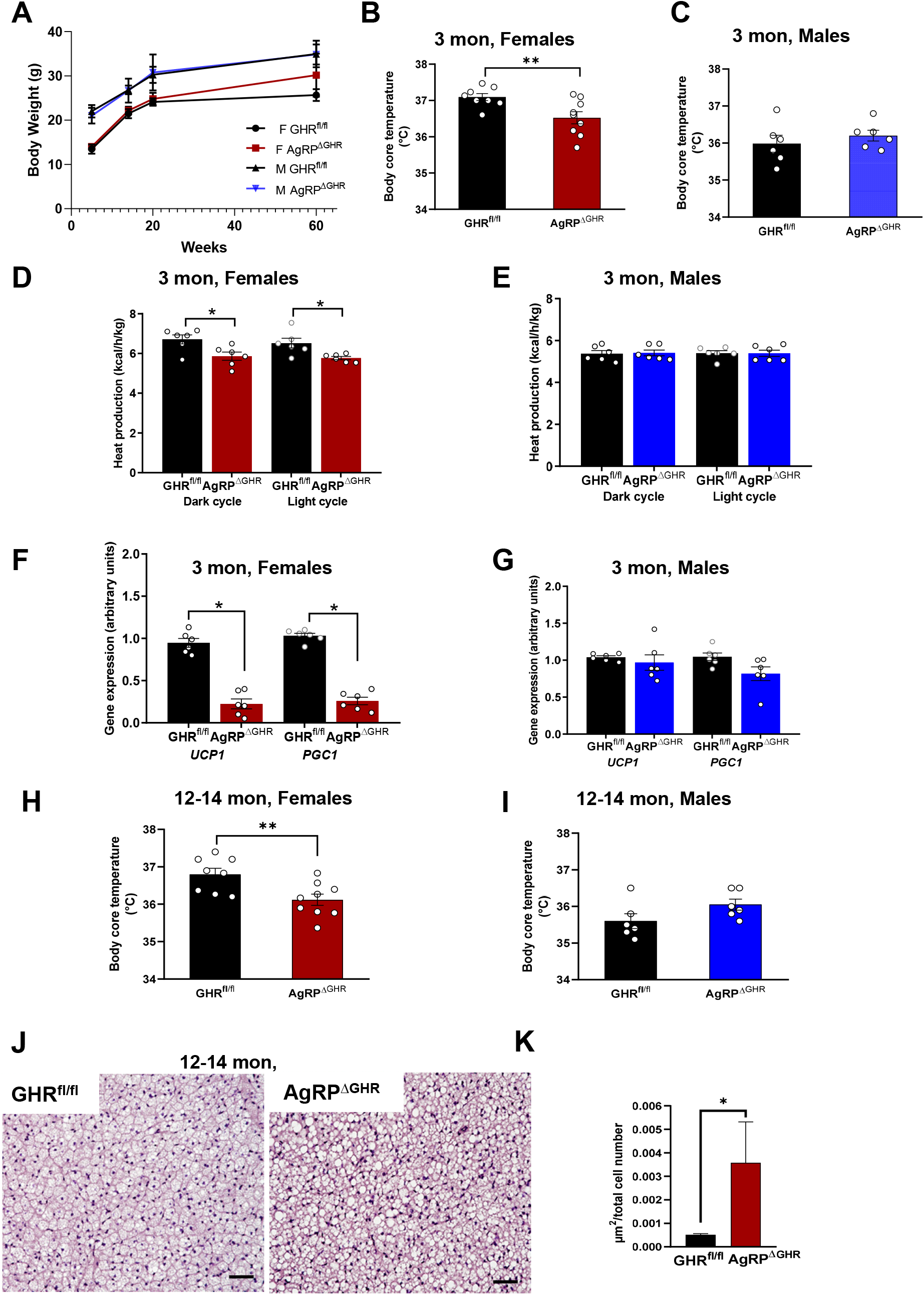
Temperature control in AgRP^ΔGHR^ mice. (A) Body weight. Body core temperature of (B) 12-week-old female and (C) male control and AgRP^ΔGHR^ mice. Energy parameters were measured in *ad libitum* control and AgRP^ΔGHR^ mice. Heat production in (D) female and (E) male mice. Gene expression of *UCP1* and *PGC1* in BAT of 12-weeks old (F) female and (G) male mice (n=6). Data are shown as mean ± SEM. *, p<0.001. Body core temperature of 12-14 months old (H) female and (I) male control and AgRP^ΔGHR^ mice (n=7-8). Representative images (J) and quantification (K) of H&E staining in BAT of 12-14 months old female mice (n=6). Data are shown as mean ± SEM. *, p<0.05. See also Supplementary Figure 1.

Interestingly, as found at a young age, 12-14 month-old female AgRP^ΔGHR^ mice exhibited decreased body core temperature compared to control. No differences were detected in body core temperature in male AgRP^ΔGHR^ mice (Figure 1H and I). Histological examination of BAT (H&E-stained sections) revealed a significant increase in lipid droplet size in brown adipocytes in young and middle-aged female AgRP^ΔGHR^ (Figure 1J, K, and Supplementary Figure 1C, D).

### Remodeling of BAT transcription in adult AgRP^ΔGHR^ female mice

To further investigate age-related aberrations in BAT we performed bulk RNA sequencing (RNA-seq) of the BAT from 14-months old female AgRP^ΔGHR^ mice and compared it to control females. Results of principal component analysis for the most variable genes and hierarchical clustering in BAT identified a clear separation for control vs AgRP^ΔGHR^ animals (Supplementary Figure 2). Volcano plot shows the main upregulated and downregulated genes in female AgRP^ΔGHR^ mice relative to the controls (Figure 2A). The complete list of regulated pathways is presented in Supplementary Tables 1–3. Of particular interest, the number of lipid metabolic pathways, including triglyceride metabolism, glycerolipid metabolism, acylglycerol biosynthesis, lipid and phospholipid biosynthesis, and isocitrate metabolism, were significantly downregulated in BAT from female AgRP^ΔGHR^ mice as compared to the controls (Figure 2B). Among the genes associated with lipid metabolic processes and glucose regulation *Apobec1, Cyp2e1, Gk, Malat1, Ankrd9, Kcnq1ot1, Slc12a2*, and *Lpl* were downregulated, while genes associated with fatty acids metabolism *Scd1* and *Scd2* were upregulated (Figure 2C).

**Figure 2:**
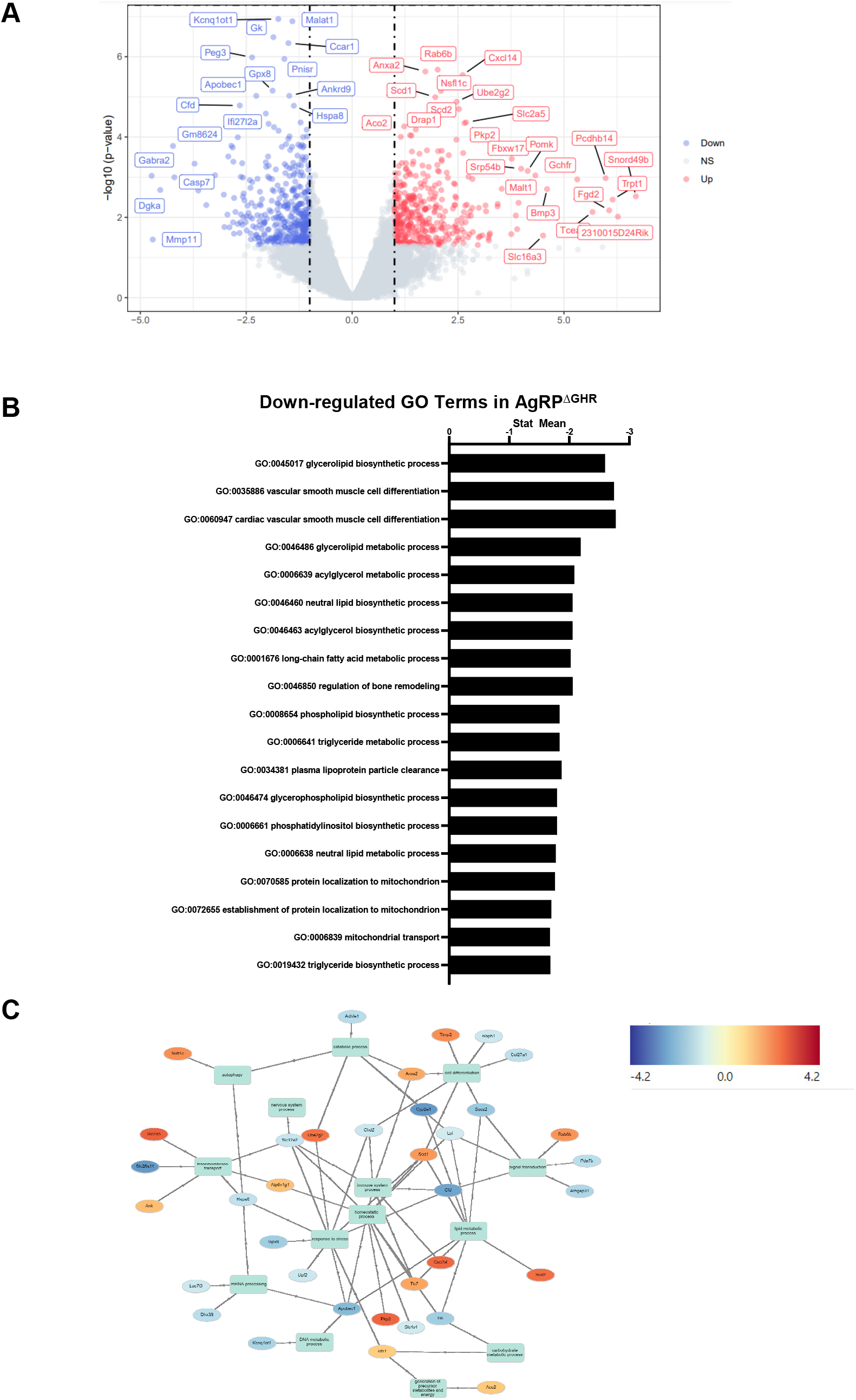
BAT transcription profile in adult AgRP^ΔGHR^ female mice. (A) Volcano plot of differential expression genes in 12-14 months old AgRP^ΔGHR^ females compared to control females. The blue, red, and grey dots represent the down-regulated, up-regulated, and unchanged genes respectively. (B) Down-regulated functions identified by GO-analysis. (C) Network analysis of differentially expressed genes associated with carbohydrate metabolic process, lipid metabolic process, immune system, homeostatic process, and stress response. Red: up-regulated genes compared to the control group; blue: down-regulated genes compared to the control group. See also Supplementary Figure 2 and Supplementary Tables for pathways analysis.

### AgRP^ΔGHR^ female mice do not adapt to changes in temperature

To further investigate the requirement of GHR in AgRP neurons for adaptation to cold or at thermoneutrality, we recorded body core temperature during the light cycle in middle-aged 12-14 months old females exposed for 3 days to 10⁰C or 30⁰C and compared the outcomes to those measured in mice housed at room temperature (22⁰C) (Figure 3A). Using this exposure paradigm, we found that female AgRP^ΔGHR^ mice maintained their low body core temperature upon exposure to 10⁰C, as compared to control (Figure 3B). Furthermore, 3 days of exposure to 30⁰C did not normalize the body core temperature of female AgRP^ΔGHR^ that was maintained consistently lower than controls (Figure 3C). Middle-aged male AgRP^ΔGHR^ mice showed adaptation to changes in temperature that was similar to their control littermates (Figure 3D, E and Supplementary Figure 3). Total body weight was unaffected (data not shown), however, the sustained cold exposure or thermoneutrality resulted in pronounced changes in BAT morphology in control animals. H&E-stained BAT from female mice housed at ambient temperature had brown adipocytes with multiple small lipid droplets, while BAT from female AgRP^ΔGHR^ mice had significantly larger single lipid droplets (Figure 3F and G). In contrast, H&E stained-BAT from cold-exposed control female mice showed much smaller brown adipocytes, and fewer lipid droplets than at 22⁰C (p<0.00001 for effect of temperature), indicating that cold exposure induced a loss of lipid droplets in the brown adipocytes (Figure 3F). We did not detect significant cold-induced morphological changes in female AgRP^ΔGHR^ mice compared to controls (Figure 3F and H). Regardless, *Ucp1* and *Pgc1* expression levels increased with cold temperature in control but not in female AgRP^ΔGHR^ mice. Importantly, the expression levels of *Ucp1* and *Pgc1* were downregulated in the female AgRP^ΔGHR^ mice housed at room temperature and maintained low regardless of the temperature change (Figure 3I). Interestingly, expression levels of *Fgf21* were also significantly elevated in response to cold exposure in control mice, supporting its activation by cold [24], while *Fgf21* levels were significantly reduced in cold but elevated at 30⁰C in female AgRP^ΔGHR^ mice, suggesting transcriptional dysregulation of *Fgf21* in BAT.

**Figure 3:**
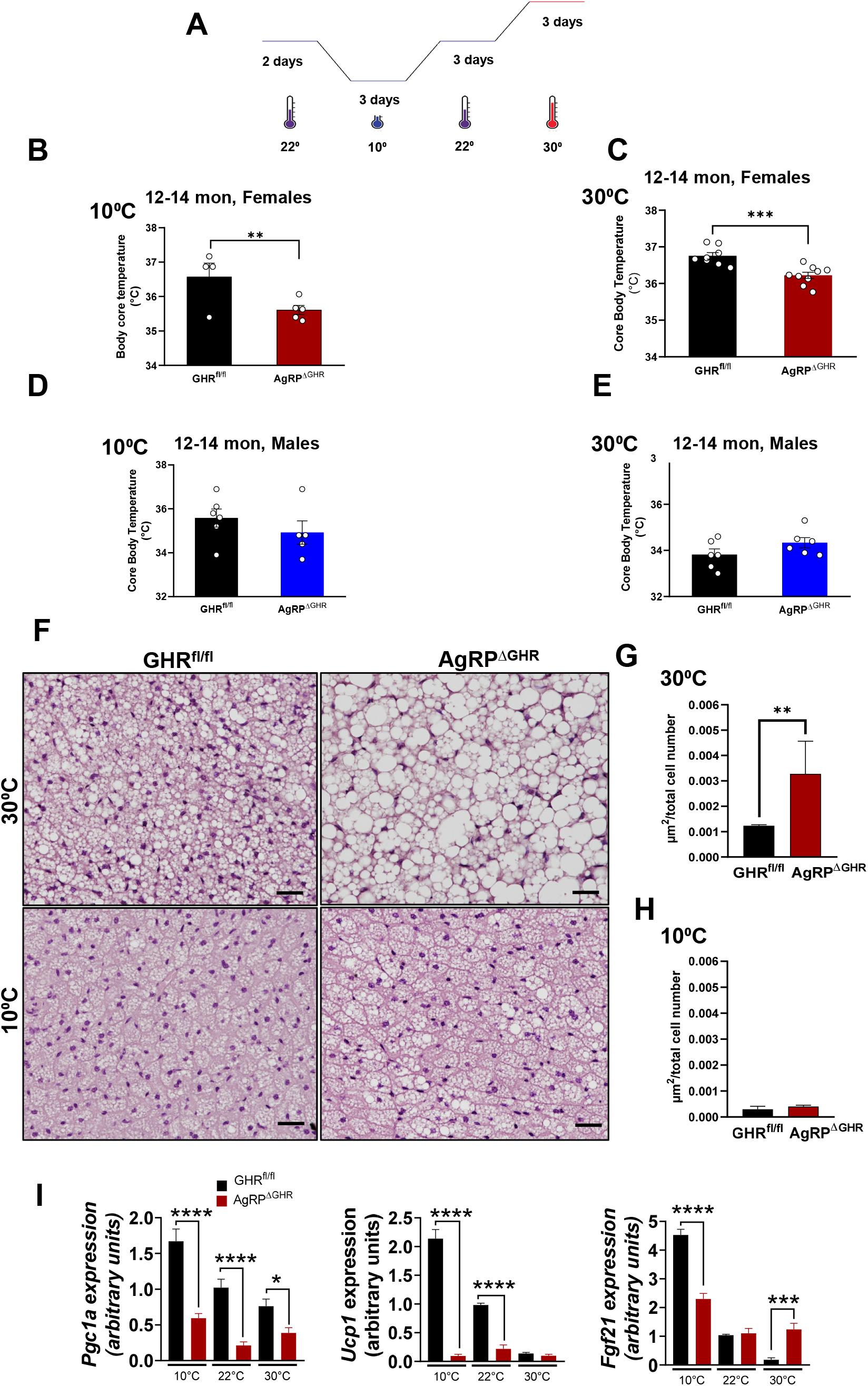
Effect of cold exposure or thermoneutrality on body core temperature and BAT gene expression. (A) 12-14 month-old control and AgRP^ΔGHR^ females and males were housed in temperature-controlled chambers set to either cold (10°C) or 30⁰C for 3 days. Body core temperature in (B) females and (D) males housed at 10°C. Body core temperature in (C) females and (E) males housed over 3 days at 30°C. n = 5–8 per group, mean ± SEM. Student’s t-test, *, p<0.05. Representative images (F) and quantification (G and H) of H&E staining in BAT of 12-14 months old female mice housed at 10°C or 30⁰C for 3 days (n=6), t-test *p<0.05, **p<0.01. See also Supplementary Figure 3 for H&E staining in BAT of 12-14 months old male mice. (I) Gene expression of *PGC1*, *UCP1*, and *Fgf21* in BAT of 12-14-month-old control and AgRP^ΔGHR^ females housed at 22°C, 10°C or 30°C as determined by qRT-PCR, mean ± SEM. Two-way ANOVA followed by Newman–Keuls test, *p<0.05, **p<0.01, ****p<0.0001 vs. 22°C.

### ARC neurons do not adapt to cold exposure in female AgRP^ΔGHR^ mice

A recent study demonstrated that mild cold exposure induces activation of cFos protein, a marker of neuronal activation in AgRP neurons [25]. We found that the total number of cFos^+^ neurons in the ARC was comparable in both control and AgRP^ΔGHR^ mice maintained at 22°C. However, the number of cFos^+^ neurons was significantly increased in response to cold exposure in control, but not in female AgRP^ΔGHR^ mice (Figures 4A and B). These data are in agreement with previous findings showing that short but not long-term exposure to thermoneutrality suppressed activation of AgRP neurons [25, 26]. The number of cFos^+^ neurons in mice of both genotypes housed at 30° was minimal and similar to those measured at 22°C (Figure 4B).

**Figure 4:**
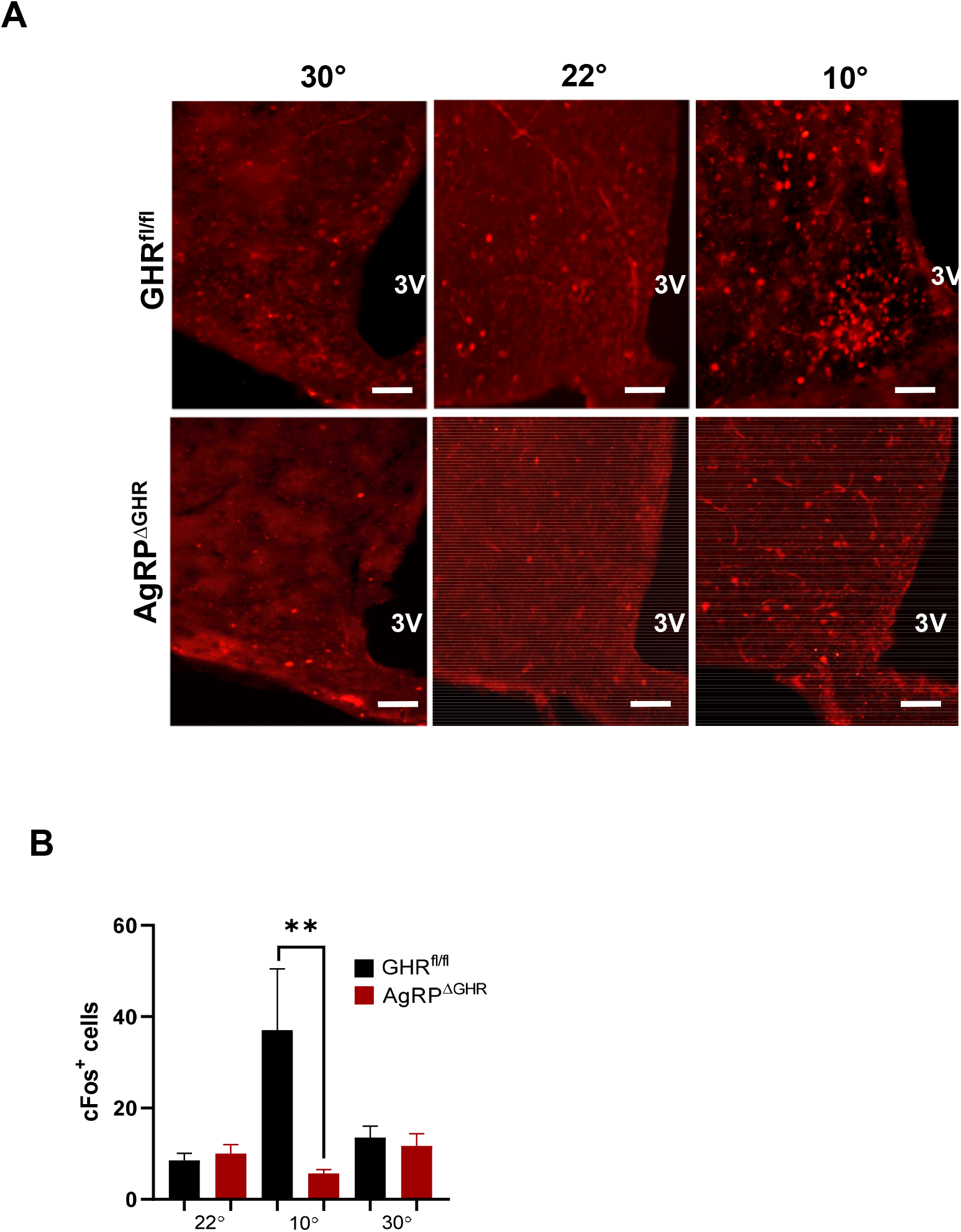
Cold exposure activates ARC neurons in control but not in AgRP^ΔGHR^ females. (A) Representative images of immunohistochemical detection of cFos (red) the arcuate nucleus (ARC) of 12-14-month-old control and AgRP^ΔGHR^ females after housing at either 10°C, 22°C, or 30°C. (B) Quantitation of Fos positive cells in the ARC. Scale bar, 100 mm, n = 3–5 group, mean ± SEM. Two-way ANOVA followed by Newman–Keuls test, **p<0.01 vs. 22°C.

### BAT transcriptome in thermoneutrality in aged female AgRP^ΔGHR^ mice

We next assessed the impact of GHR deletion from AgRP neurons on BAT transcriptome in animals adapted to thermoneutrality for 3 days as compared to animals housed at room temperature. A total of 51 genes were changed in BAT by 30⁰C in female AgRP^ΔGHR^ mice, compared to 101 genes in control mice (false discovery rate [FDR] < 0.05, Figure 5A) suggesting different mechanisms of adaptation. The regulated pathways for control and AgRP^ΔGHR^ females are shown in Supplementary Material. Female AgRP^ΔGHR^ mice were less responsive to thermoneutrality, with reduced changes in pathways when adapted from 22⁰C to 30⁰C (Figure 5B). Surprisingly, we identified several lipid metabolic pathways that were uniquely downregulated in control animals in adaptation to thermoneutrality and in AgRP^ΔGHR^ compared to control mice at 22⁰C (Figure 5B). Specifically, lipid biosynthesis, glycerolipid metabolism, and lipid oxidation pathways were among those uniquely downregulated in control animals in adaptation from 22⁰C to 30⁰C. In the AgRP^ΔGHR^ females, these pathways were already downregulated at 22⁰C and did not change with adaptation from 22⁰C to 30⁰C (Figure 5B and C). Among the common genes downregulated in both control and AgRP^ΔGHR^ mice by adaptation to thermoneutrality were metabolic genes *Gk, Dio2, Ucp3, Ucp1, and Ankrd9* (Figure 5D). Importantly, the genes associated with lipid accumulation and glucose regulation (*Malat1, Ankrd9, Kcnq1ot1, Slc12a2, Peg3)* which were downregulated only in control females in adaptation to 30⁰C, were also downregulated in female AgRP^ΔGHR^ mice at 22⁰C compared to control mice (Figure 5E and F). These genes were not changed in the female AgRP^ΔGHR^ mice in adaptation to 30⁰C (Figure 5F), suggesting the role of GHR in AgRP neurons for the adaptation of BAT lipid and glucose metabolism to environmental temperatures.

**Figure 5:**
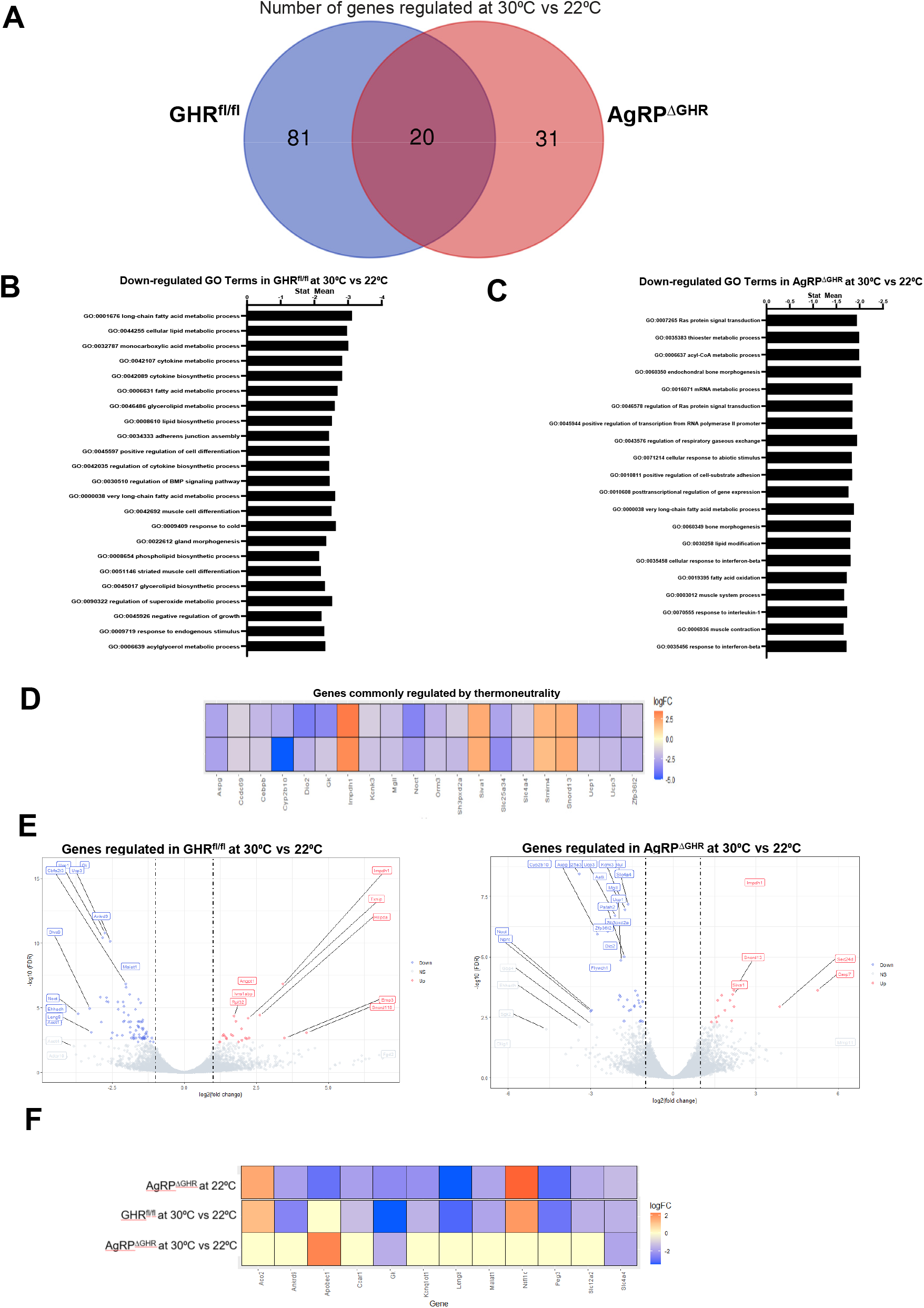
Differentially expressed genes in AgRP^ΔGHR^ females exposed to thermoneutrality. (A) The Venn diagram depicts the number of overlapping genes and differentially expressed genes between 12-14 month-old control and AgRP^ΔGHR^ females (left) at 30°C versus 22°C. (B) Down-regulated functions were identified by GO-analysis in control females at 30°C versus 22°C. (C) Down-regulated functions were identified by GO-analysis in AgRP^ΔGHR^ females at 30°C versus 22°C. (D) Heatmap illustrating the relative expression of overlapping genes by 30°C. (E) Genes regulated only in control females (right) or only in AgRP^ΔGHR^ females (left) at 30⁰C vs 22⁰C. (F) Heatmap illustrating the differentially expressed genes in AgRP^ΔGHR^ females at 22⁰C vs control, and the relative expression of unique genes at 30⁰C vs 22⁰C in control and AgRP^ΔGHR^ females.

## Discussion

We identified a unique role for GHR signaling in hypothalamic AgRP neurons in controlling thermal adaptation. Using previously characterized AgRP-specific GHR knockout mice [18], we show that GHR signaling in AgRP neurons regulates body core temperature in females but not in male mice. Importantly, middle-aged female AgRP^ΔGHR^ mice show impaired adaptation to cold or thermoneutrality with increased BAT lipids accumulation and aberrant transcriptomic signatures. Specifically, female AgRP^ΔGHR^ mice exhibited transcriptomic signatures of downregulated lipid metabolic genes that are similar to those of control females adapting to temperature change. This indicates that GHR signaling mediates the response to thermoneutrality in the AgRP neurons in a sex-specific manner.

The ARC is a major site for the integration of multiple nutritional and hormonal signals, which are central to the modulation of energy balance and temperature homeostasis [27]. Evidence for the importance of GH-responsive neurons in the hypothalamus in energy homeostasis and nutrient deprivation was previously reported [18, 28–30]. Moreover, the orexigenic effect of GH signaling is possibly mediated by AgRP neurons, since most of the AgRP/NPY neurons express GHR, and AgRP-specific GHR-knockout mice are unable to adapt to food restriction and maintained a higher energy expenditure, having increased weight loss during food restriction compared to the control mice [17]. In support, our earlier study with young male and female AgRP^ΔGHR^ mice do not show changes in body weight, body composition, food intake, or glucose homeostasis [18]. Here we show that despite similar body weights and activity levels, the obligatory energy expenditure required for basal activity is reduced in young AgRP^ΔGHR^ female, but not male mice and is associated with lower body core temperature, reduced heat production, increased BAT adipocytes size, and reduced BAT expression of thermogenic genes. During the aging process, such metabolic imbalance led to a slight increase in middle-aged females’ body weight, accompanied by an inability to properly respond to changes in environmental temperatures. Given a significant increase in BAT lipids accumulation, the duration of eating/fasting patterns may be different in AgRP^ΔGHR^ female mice since AgRP neurons regulate complex behavioral and physiological feeding changes [31, 32]. It is reasonable to hypothesize that age-associated decline in GH signaling in the AgRP neurons is responsible for or importantly contributes to the impairment in BAT thermogenic capacity occurring with age in female mice [33].

Recent work has established that in addition to negative feedback regulation of hormonal signals to nutrient intake and energy metabolism, AgRP neuron activity rapidly increases following exposure to a mild cold environment and that the activity of these neurons at thermoneutrality is lower [25]. Moreover, exposure to a warm environment was shown to suppress AgRP activity in 10-day-old pups [26]. A recent study demonstrated that the control of BAT thermogenesis by AgRP neurons is independent of environmental temperature and activation of thermoregulation since specific activation of AgRP neurons suppresses BAT thermogenic activity [34]. Specifically, in mice maintained in the thermoneutral zone of 30°C, the effects of AgRP neuronal activation on BAT temperature, energy expenditure, and locomotor activity were significantly reduced. Moreover, during cold exposure, chemogenetic activation of AgRP neurons reduced BAT temperature to a similar extent as at ambient temperature, providing evidence for the AgRP-BAT circuit independent of environmental temperature and cold-induced thermoregulation. This effect was mediated via hypothalamic mTOR complex-1 (mTORC1) signaling [34].

Middle-aged AgRP^ΔGHR^ female mice exhibit reduced body core temperature regardless of environmental temperature (22°C, 10°C, or 30°C), suggesting that GHR signaling in the AgRP neurons, can sense energy availability and AgRP activity related to AgRP-BAT circuit. mTORC1 activity is indeed lower in multiple organs of Snell dwarf and global GHRKO mice [35], suggesting a requirement of this mechanism and properly regulated GHR signaling in AgRP neurons for thermogenic action. Indeed, we found down regulation of mTOR signaling in control BAT but not in BAT of AgRP^ΔGHR^ females. Further insights on the regulation of the mTORC1 pathway in the AgRP neurons lacking GHR signaling would be informative to establish the role of this circuit in the regulation of body core temperature in aging.

Our new data provide critical insight into the sexual dimorphism of GHR signals in AgRP neurons in thermoregulation. While unexpected, a similar sex-specific phenotype was shown in females lacking corticotropin-releasing factor receptor type 1 (CRFR1) in AgRP neurons [11]. Only female mice selectively lacking CRFR1 in AgRP neurons exhibited reduced heat production, and lower body temperature, followed by a maladaptive thermogenic response to cold. In the ARC, the GHR gene is co-expressed with CRFR in the same cluster of AgRP neurons [36], suggesting a sex-specific role of this subpopulation of AgRP neurons in energy homeostasis and regulation of body core temperature. Additionally, a sex-specific phenotype was observed in females lacking leptin receptors in POMC neurons or AgRP neurons, which, as in AgRP^ΔGHR^ and AgRP^ΔCRFR1^ females, exhibited reduced heat production with unaltered food intake [37, 38]. These sex-specific differences can be attributed to hypothalamic estrogen signals [39]. There is some evidence that estrogen modulates GH action independent of secretion [40]. While estrogen receptor is not expressed by AgRP neurons [41], estrogen can directly affect GHR expression and signaling [40]. However, it is important to note that within each sex, the levels of GH, were unaltered in AgRP^ΔGHR^ mice [17, 18]. Our new studies thus provide a critical view of the sex-specific role of GHR signaling in AgRP neurons in thermoregulation with particular importance for aged females.

Transcriptomic analysis of BAT demonstrated that genes associated with lipid metabolic processes and lipids accumulation were downregulated in AgRP^ΔGHR^ compared to control. Moreover, these were the same genes downregulated by thermoneutrality in control mice but not in the AgRP^ΔGHR^ animals. The divergent response of BAT in the AgRP^ΔGHR^ females was evident by downregulation in key genes associated with lipid accumulation and glucose regulation such as *Malat1, Ankrd9, Kcnq1ot1, Slc12a2, and Peg3.* These genes were not changed in the AgRP^ΔGHR^ females during adaptation to 30⁰C. Some of these genes, such as *Malat1* were shown to promote hepatic steatosis and insulin resistance by modulating lipid accumulation through SREBP-1c [42], while *Ankrd9*, *Kcnq1ot1*, *Peg3* are involved in intracellular lipid accumulation, and lipogenesis [43–45], and *Slc12a2* plays a role in insulin secretion and glucose-stimulated plasma membrane depolarization [46]. This selective alteration in the genes is regulated by the acclimation to a thermoneutral environment in middle-age control females resembles the changes detected in the AgRP^ΔGHR^ mice at 22⁰C. This suggests that these animals were capable of maintaining the basal metabolic rate of low euthermia thought their life span regardless of the temperature exposure. Interestingly, genes associated with fatty acids metabolism *Scd1* and *Scd2* were upregulated in the AgRP^ΔGHR^ females indicating shifts in metabolic homeostasis and substrate utilization [47]. Interestingly, SCD1 deficiency stimulates basal thermogenesis through the upregulation of the beta-3-AR-mediated pathway and an increase in lipolysis and fatty acid oxidation in BAT [48].

A relationship between GH and thermoregulation was previously suggested. Central infusion of GH into the hypothalamus leads to an increase in sympathetic nerve activity [49, 50]. Furthermore, the inability of whole-body GHRKO mice to respond to cold exposure or β-adrenergic receptor agonist treatment suggested that GH plays an important role in thermoregulation [51]. Previous studies using whole-body GHRKO and GH deficient Ames dwarf mice, showed an increased amount of BAT [14, 52], increased expression of thermogenic genes in BAT, and thermogenic activation (‘browning”) of subcutaneous white adipose tissue (WAT), which can stimulate thermogenesis [53]. Intriguingly, similarly to AgRP^ΔGHR^ females, body core temperature was reduced in these mutants [54, 55] despite increased thermogenesis and metabolic rate. Reduced body temperature has been associated with extended longevity [56]. Additionally, GH-deficient mice exposed to thermoneutrality had reduced expression of genes associated with lipid metabolism and energy expenditure, and morphological changes in BAT similar to AgRP^ΔGHR^ females [22]. More than 1500 genes are regulated in BAT at room temperature or thermoneutrality in global GHRKO mice [22]. When GHRKO mice were exposed to thermoneutral conditions, a greater number of genes were affected in BAT from GHRKO than control mice, with low overlap between groups, indicating a divergent response. Similar to the global GHRKO mice, AgRP^ΔGHR^ mice, showed little overlap in genes when compared to the responses of control mice to thermoneutrality. Nevertheless, 114 genes were commonly regulated in response to thermoneutrality by global GHRKO and AgRP^ΔGHR^. Among the pathways regulated by these common genes, lipid metabolic process, fatty acid metabolic process, response to cold, and brown fat cell differentiation were the top regulated GO terms [22]. Similarly to AgRP^ΔGHR^ mice, *Ankrd9* was upregulated in GHRKO mice at 22⁰C but downregulated in GHRKO mice exposed to thermoneutral conditions [22]. On the other hand, *Peg3* was upregulated in GHRKO compared to WT mice at 22⁰C, while *Slc12a2* and *Scd1* were downregulated in GHRKO mice exposed to thermoneutrality [22], reflecting the differences between both models. Additional work will be required to define the common and divergent responses between these models in the regulation of body core temperature and energy homeostasis.

Aging is associated with an attenuated physiological ability to maintain body core temperature and the risk of heat-related illness in these individuals is elevated [57]. The role of sex and reproductive hormones in thermoregulation is complex and depends on the situation, overall health, and age [58]. Our data provide evidence that GHR signaling in the AgRP neurons mediates the response to thermoneutrality, controlling lifetime steady state temperature in female animals. Future studies would be required to define the impact of low body core temperature in AgRP^ΔGHR^ female mice on longevity and metabolic health in aging.

## Abbreviations

CNS: central nervous system
GH: growth hormone
GHR: growth hormone receptor
POMC: proopiomelanocortin
ARC: arcuate nucleus of the hypothalamus
DMH: dorsomedial hypothalamic nucleus
LHA: lateral hypothalamus
PVH: paraventricular hypothalamic nucleus
GHRH: growth hormone releasing hormone
AgRP: agouti-related peptide
BAT: brown adipose tissue

## Author contribution

LS, JBML, LKD, carried out the research and reviewed the manuscript. MK and LK assisted in data collection and experimental design. JJK contributed the GHR flox mice. AS and AB analyzed the data and reviewed the manuscript. MS designed the study, analyzed the data, wrote the manuscript, and is responsible for the integrity of this work. All authors approved the final version of the manuscript.

## Data Availability Statement

The data that support the findings of this study are available from the corresponding author upon reasonable request.

## SUPPLEMENTARY MATERIAL

### Supplementary Figures

**Supplementary Figure 1:**
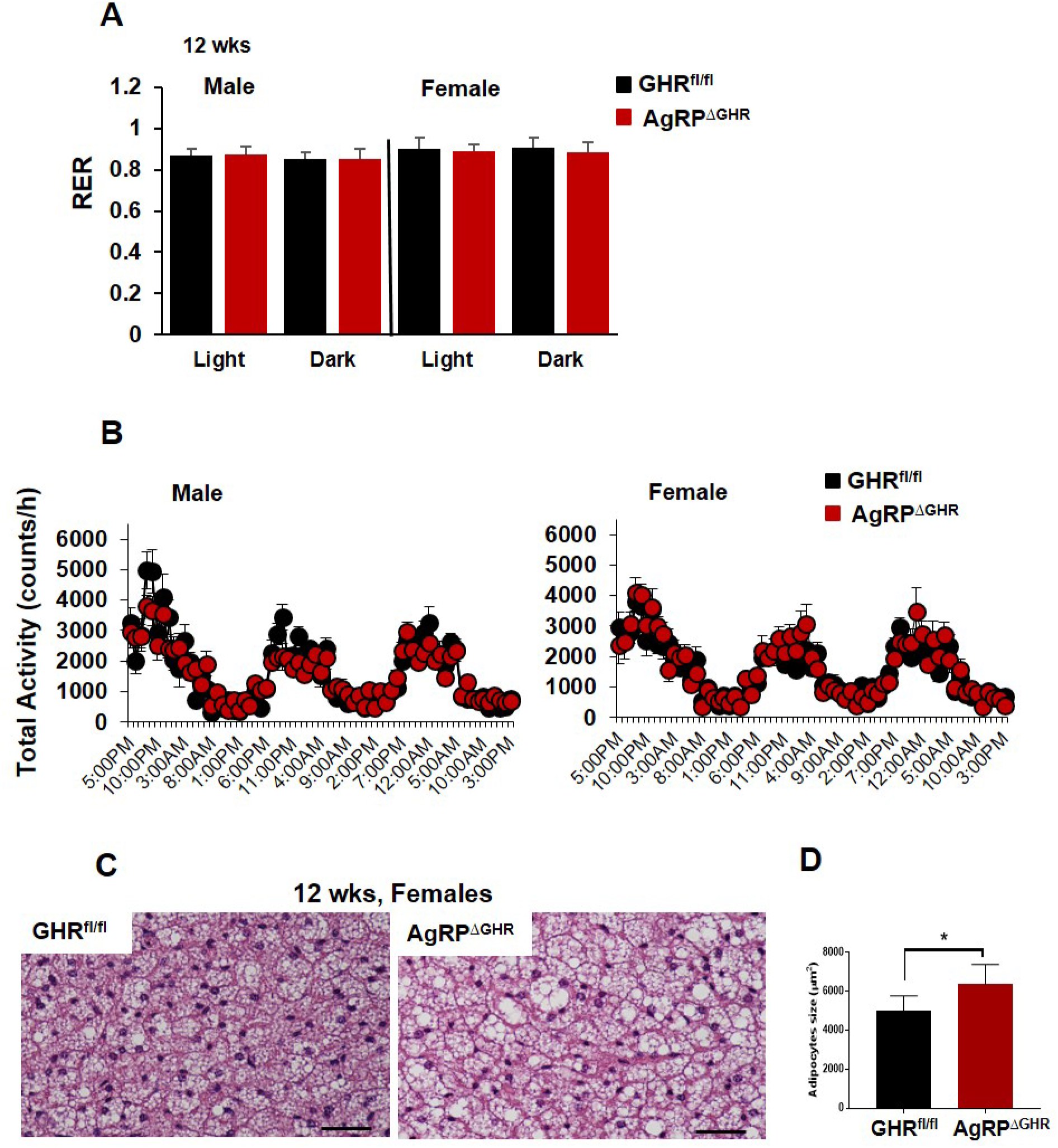
Energy homeostasis parameters in 12-week-old control and AgRP^ΔGHR^ male and female mice. (A) RER (B) Total locomotor activity. (C) Representative H&E staining of brown adipose tissue (BAT) of control (left panel) and AgRP^ΔGHR^ female mice (right panel) (n=6). Data are shown as mean ± SEM. *, p<0.005. Data relates to Figure 1

**Supplementary Figure 2:**
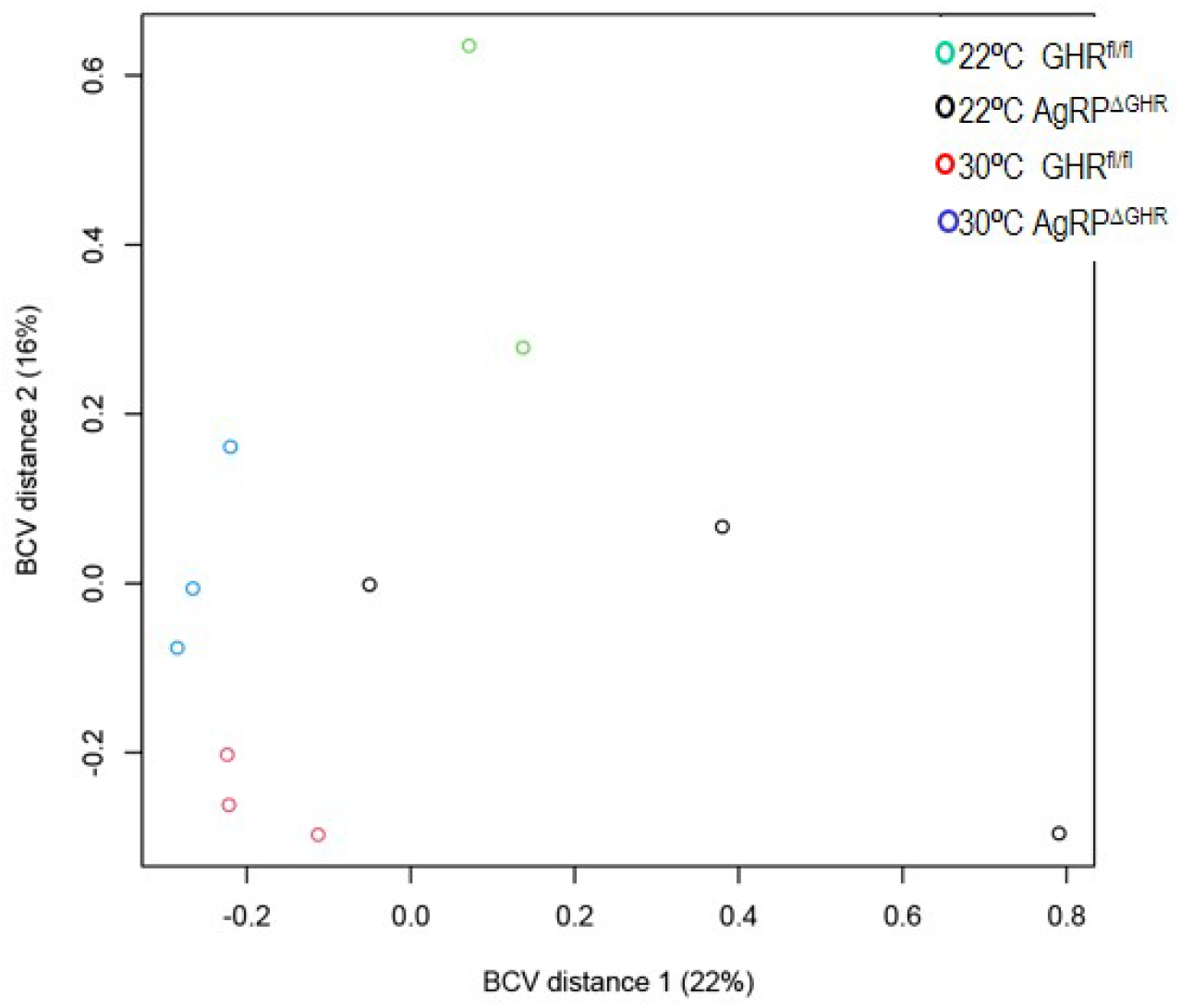
Principal components analysis of the most variable expressed mRNAs in brown adipose tissue (BAT). Control and AgRP^ΔGHR^ female mice at 22⁰C and at 30 ⁰C

**Supplementary Figure 3:**
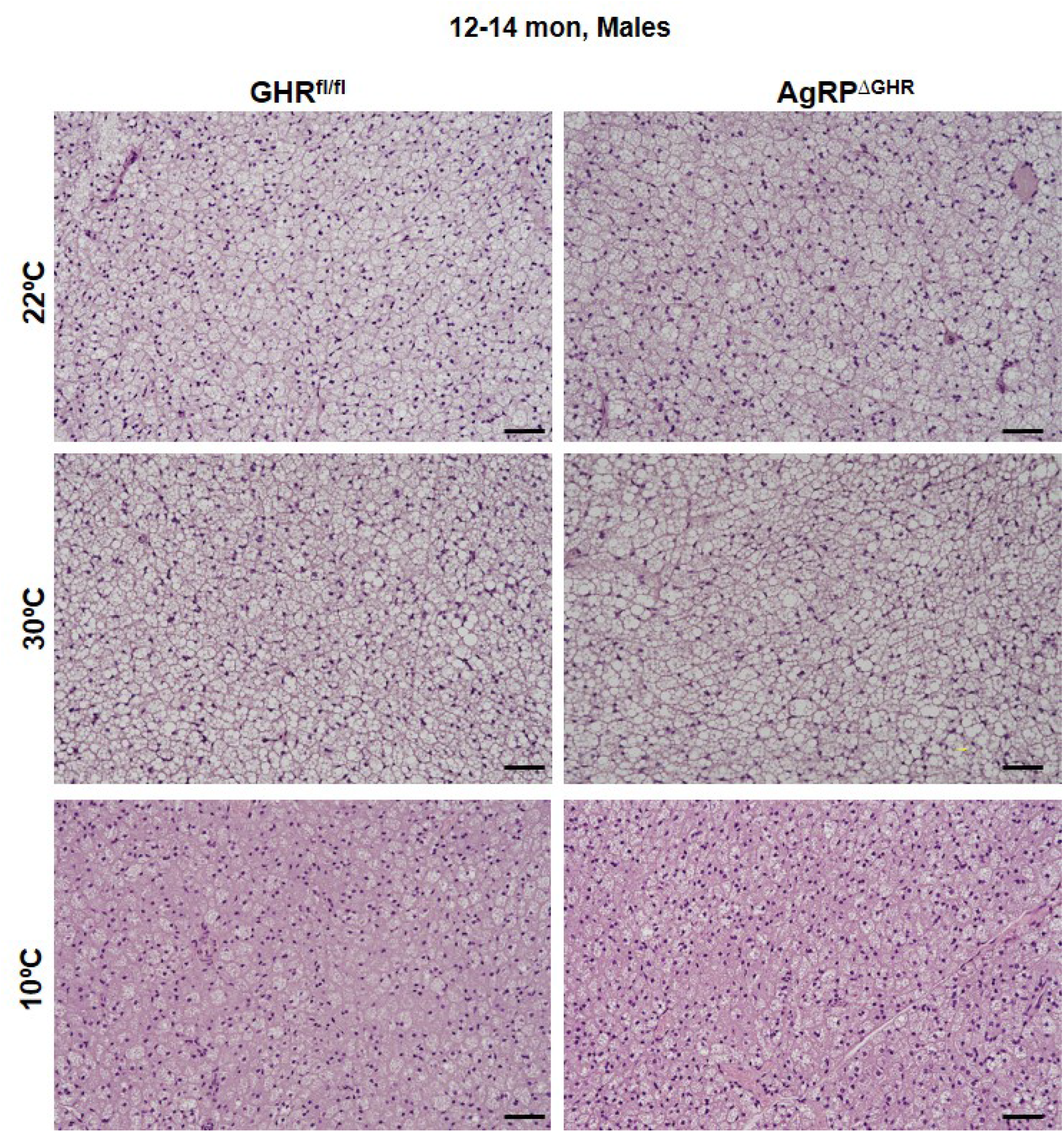
Energy homeostasis parameters in 12-week-old control and AgRP^ΔGHR^ male and female mice. (A) RER (B) Total locomotor activity. (C) Representative H&E staining of brown adipose tissue (BAT) of control (left panel) and AgRP^ΔGHR^ female mice (right panel) (n=6). Data are shown as mean ± SEM. *, p<0.005. Data relates to Figure 3

### Supplemental tables

**Suppl. Table 1.**
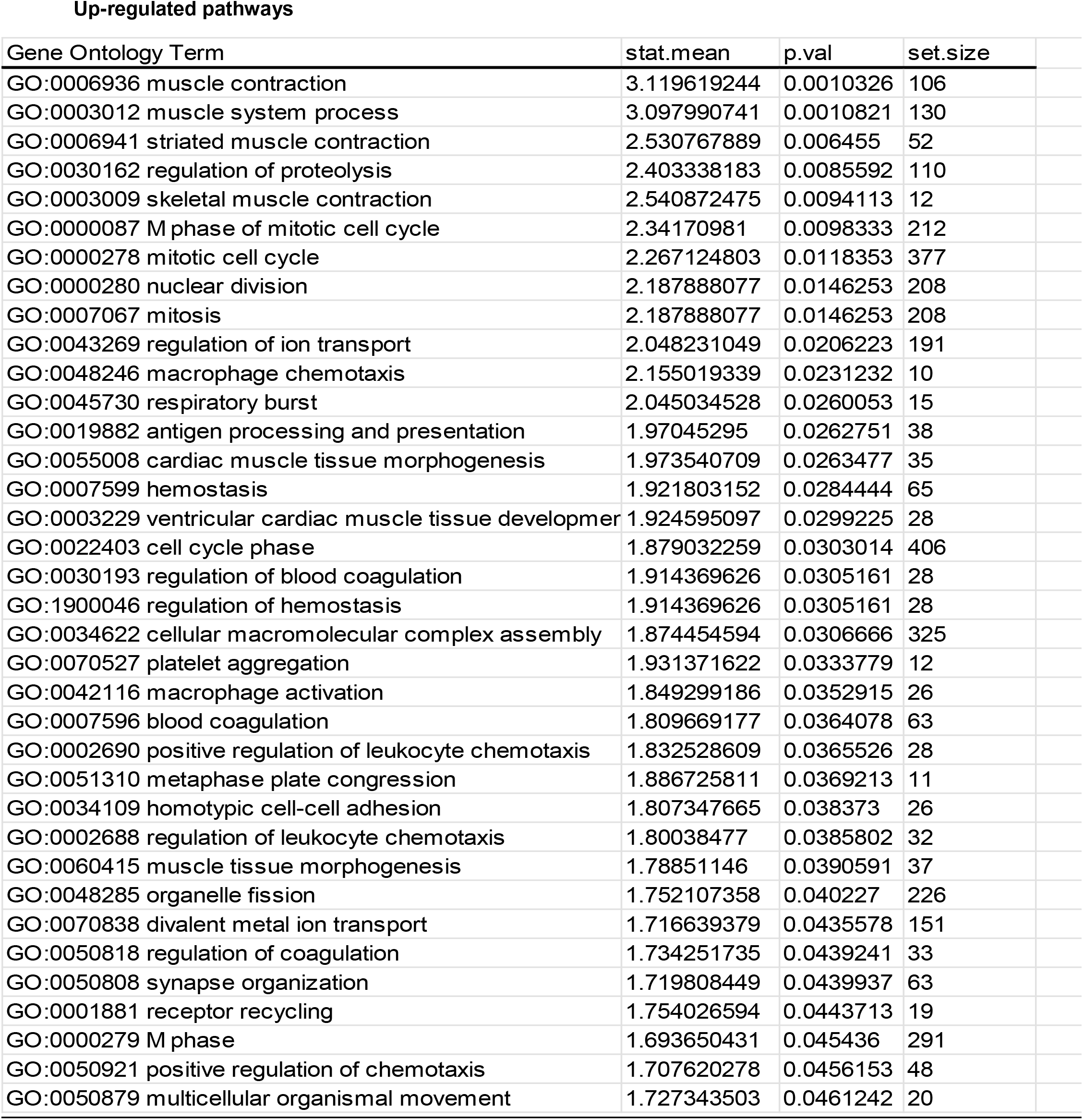
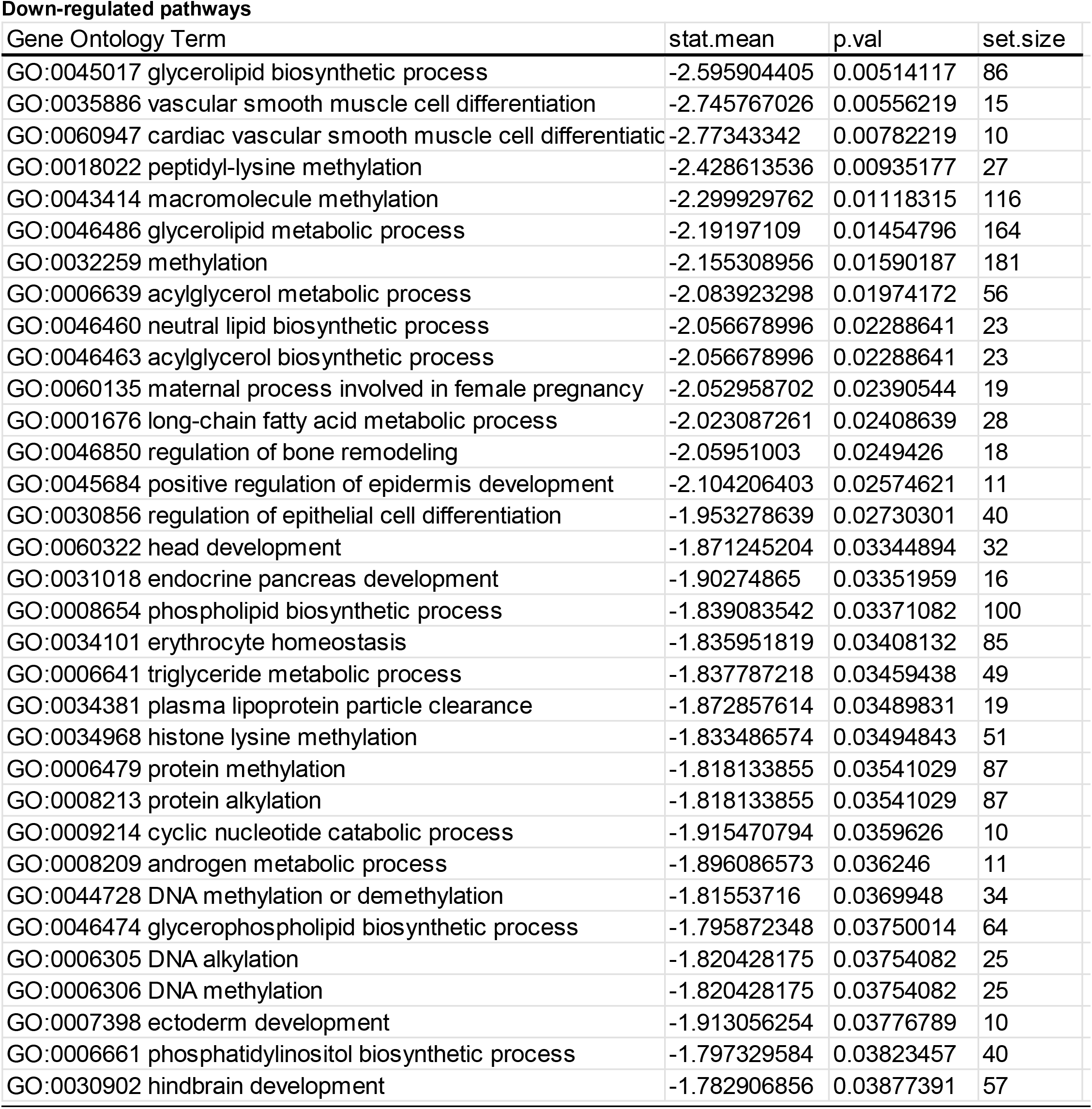
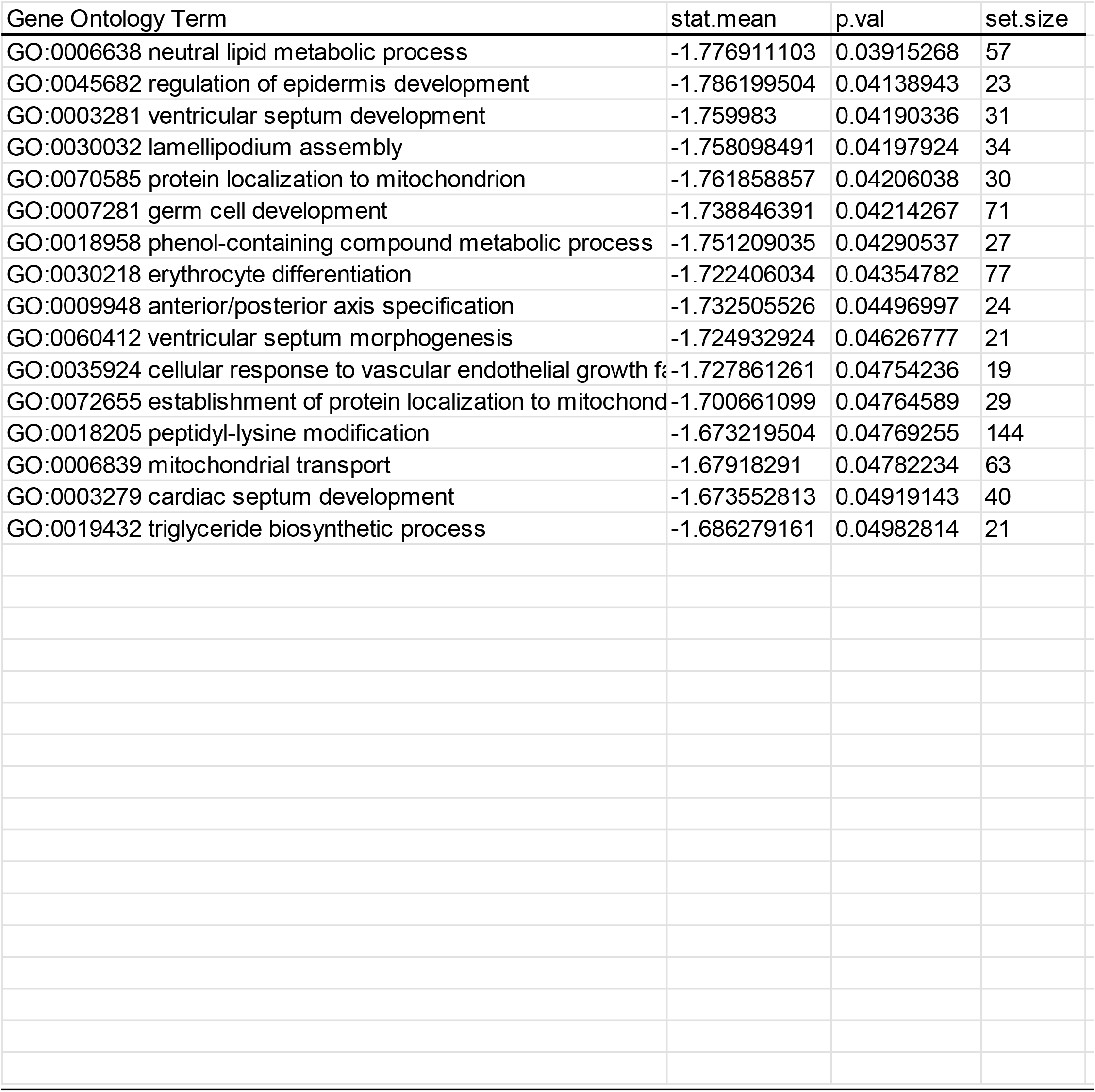
Enrichment of gene ontology terms in BAT from AgRP^ΔGHR^ mice at 22°

**Suppl. Table 2.**
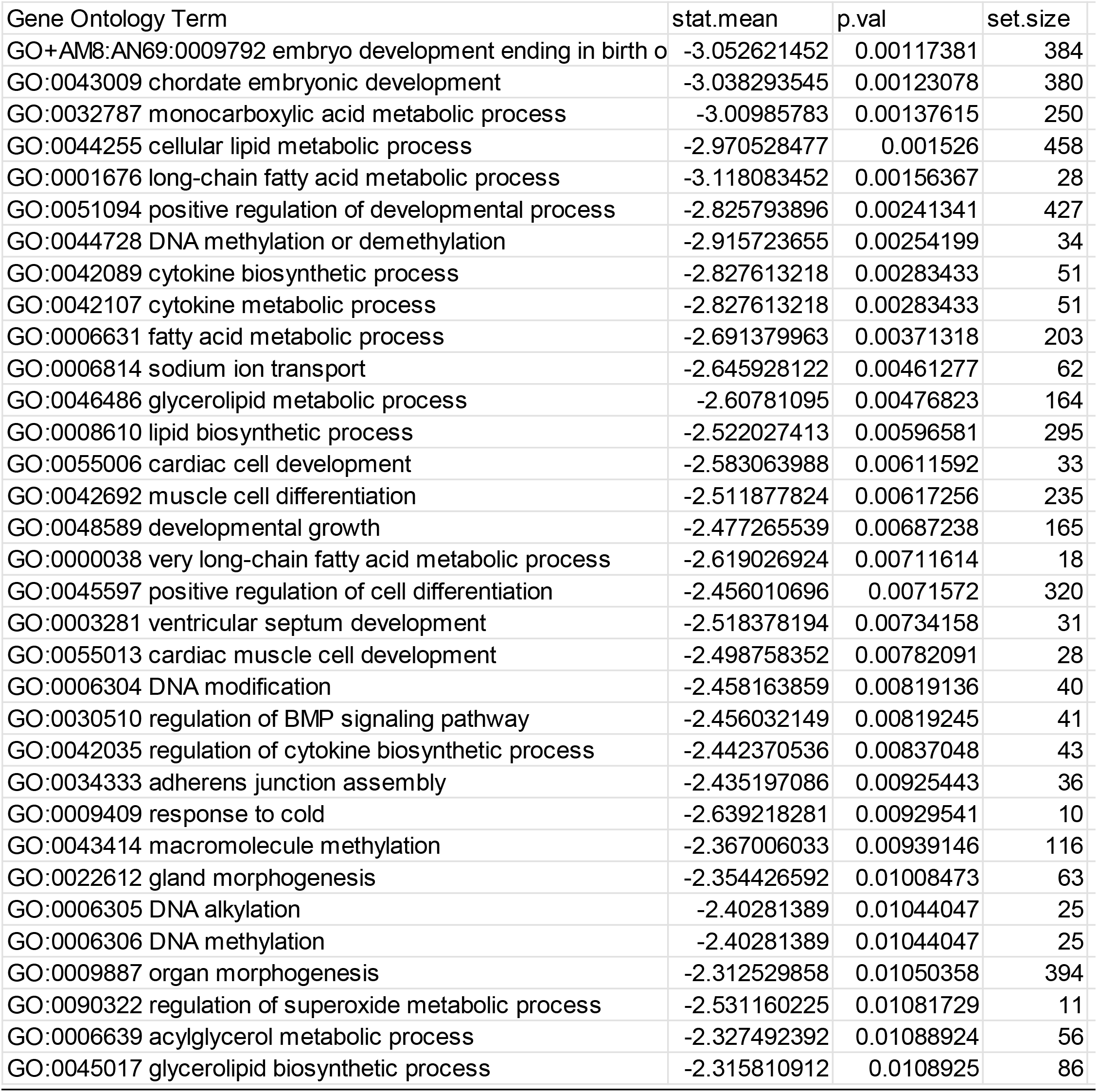
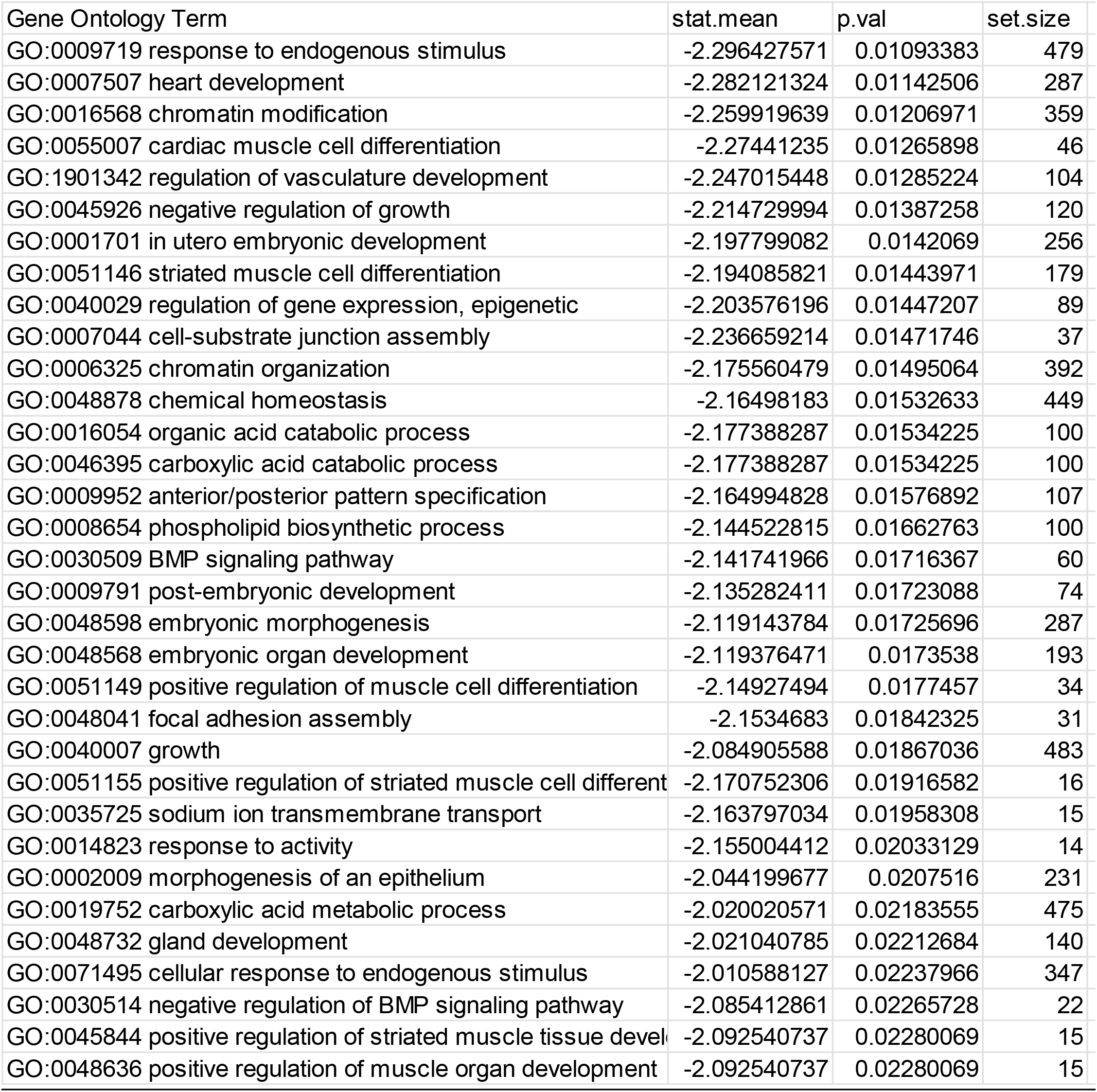
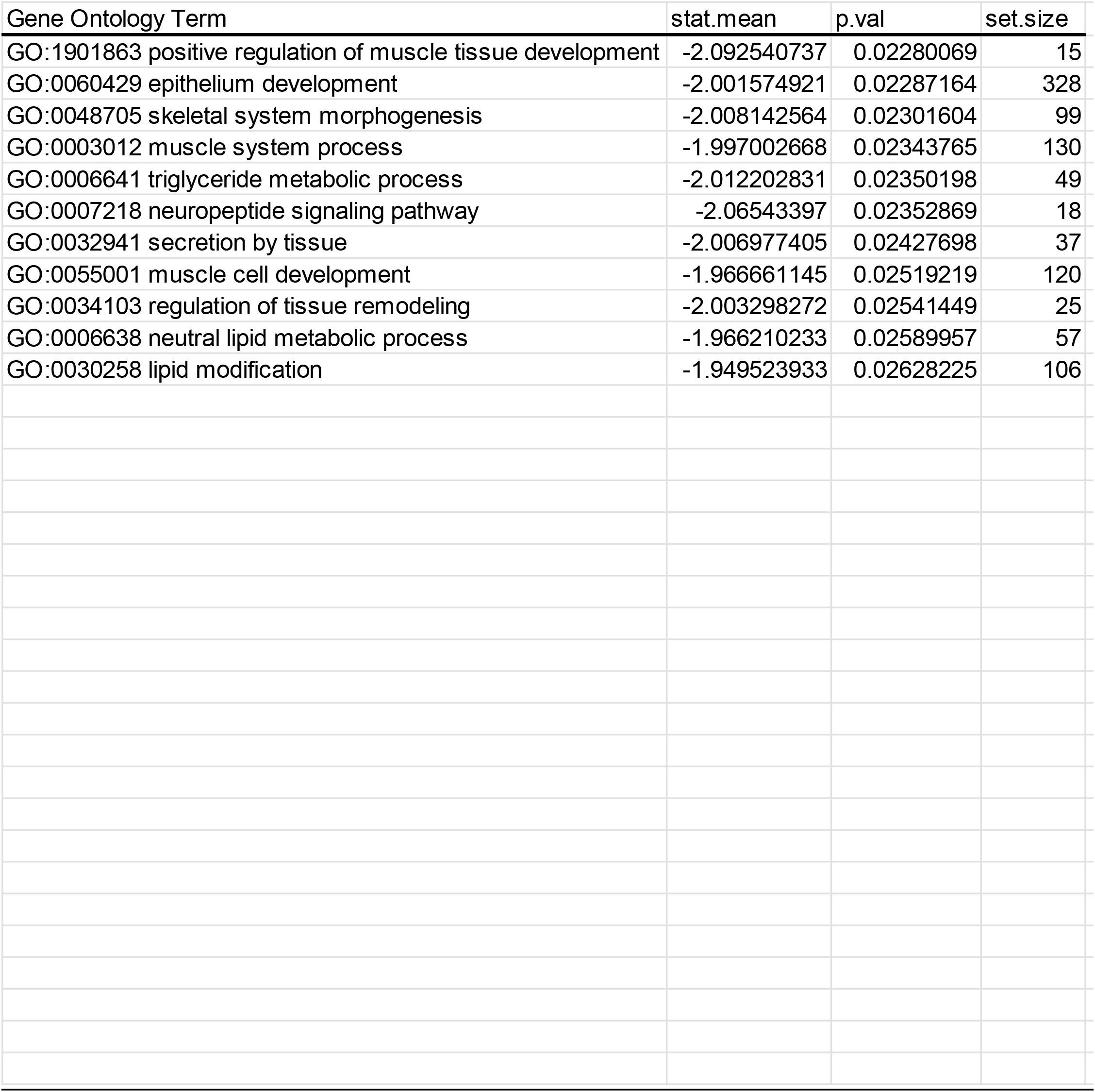
Enrichment of gene ontology terms in BAT from GHR^fl/fl^ mice at 30° vs 22°

**Suppl. Table 3.** Enrichment of gene ontology terms in BAT from AgRP^ΔGHR^ mice at 30°vs 22°

| Gene Ontology Term | stat.mean | p.val | set.size |
| --- | --- | --- | --- |
| GO:0006325 chromatin organization | -2.272750379 | 0.01165907 | 392 |
| GO:0016568 chromatin modification | -2.149459978 | 0.0159719 | 359 |
| GO:0051056 regulation of small GTPase mediated signal trans | -2.143727659 | 0.01629126 | 230 |
| GO:0007266 Rho protein signal transduction | -2.04966631 | 0.02092573 | 92 |
| GO:0006882 cellular zinc ion homeostasis | -2.175643554 | 0.02205032 | 10 |
| GO:0055069 zinc ion homeostasis | -2.175643554 | 0.02205032 | 10 |
| GO:0060350 endochondral bone morphogenesis | -2.030942362 | 0.02431471 | 22 |
| GO:0006637 acyl-CoA metabolic process | -1.989269522 | 0.02523296 | 41 |
| GO:0035383 thioester metabolic process | -1.989269522 | 0.02523296 | 41 |
| GO:0007265 Ras protein signal transduction | -1.940530929 | 0.02658181 | 167 |
| GO:0051276 chromosome organization | -1.867425485 | 0.03107361 | 485 |
| GO:0046578 regulation of Ras protein signal transduction | -1.84903222 | 0.03258235 | 209 |
| GO:0016071 mRNA metabolic process | -1.846848203 | 0.03261362 | 328 |
| GO:0045944 positive regulation of transcription from RNA polymerase | -1.841977075 | 0.0328983 | 468 |
| GO:0051568 histone H3-K4 methylation | -1.882024158 | 0.03310752 | 28 |
| GO:0043576 regulation of respiratory gaseous exchange | -1.945530803 | 0.0337513 | 10 |
| GO:0071214 cellular response to abiotic stimulus | -1.831258696 | 0.03452422 | 77 |
| GO:0010811 positive regulation of cell-substrate adhesion | -1.841054414 | 0.03462512 | 42 |
| GO:0000038 very long-chain fatty acid metabolic process | -1.875335906 | 0.03527631 | 18 |
| GO:0051310 metaphase plate congression | -1.901821235 | 0.03587092 | 11 |
| GO:0030258 lipid modification | -1.795879951 | 0.03697851 | 106 |
| GO:0060349 bone morphogenesis | -1.811164171 | 0.037039 | 39 |
| GO:0006814 sodium ion transport | -1.801676724 | 0.03711173 | 62 |
| GO:0040029 regulation of gene expression, epigenetic | -1.792304896 | 0.03740501 | 89 |
| GO:0001881 receptor recycling | -1.834722711 | 0.03747828 | 19 |
| GO:0010608 posttranscriptional regulation of gene expression | -1.762124219 | 0.03931694 | 271 |
| GO:0035458 cellular response to interferon-beta | -1.807395593 | 0.03977913 | 18 |
| GO:0019395 fatty acid oxidation | -1.724654221 | 0.04365222 | 58 |
| GO:0000375 RNA splicing, via transesterification reactions | -1.713839884 | 0.04431454 | 79 |
| GO:0070555 response to interleukin-1 | -1.732001629 | 0.04496747 | 28 |
| GO:0000377 RNA splicing, via transesterification reactions with | -1.69005115 | 0.04656134 | 78 |
| GO:0000398 mRNA splicing, via spliceosome | -1.69005115 | 0.04656134 | 78 |
| GO:0035456 response to interferon-beta | -1.716756352 | 0.0468849 | 21 |

